# Detrimental effects of atomoxetine on visual signal detection in rats: Comparison with ADHD psychomotor stimulant drugs

**DOI:** 10.64898/2026.03.06.710082

**Authors:** Livia J.F. Wilod Versprille, Koji Yano, Anders Petersen, Jeffrey W. Dalley, Trevor W. Robbins

## Abstract

**Rationale:** Attention-deficit/hyperactivity disorder (ADHD) is associated with executive dysfunction involving inattention and impulsivity, with evidence of disrupted functional expression of the dopamine and noradrenaline transporters.

**Objective:** We investigated the dose-dependent modulation of anti-ADHD drugs on selective and sustained visual attention in low-, mid-and high-attention phenotypes. Two mathematical approaches, signal detection theory and theory of visual attention were applied to further characterise the effects and mechanisms.

**Methods:** Rats were trained to detect and respond to the presence or absence of a visual target to obtain food reward on a signal detection task. After attentional performance stabilised, the indirect catecholamine agonist, d-amphetamine (0.1; 0.2; 0.4 mg/kg), the dopamine (DA) and noradrenaline (NA) reuptake inhibitor methylphenidate (0.3; 1; 3 mg/kg), and the NA reuptake inhibitor atomoxetine (0.1; 0.3; 1 mg/kg), were administered systemically.

**Results:** Low-dose d-amphetamine produced baseline-dependent effects on attention, improving target discrimination only in rats with lower attentive performance, whereas methylphenidate did not significantly improve attention but increased guessing. In contrast, low-dose atomoxetine selectively impaired attention in low-attentive subjects, whereas high-dose atomoxetine generally impaired discrimination performance. All three drugs had expected effects on motor response output.

**Conclusions:** As well as demonstrating baseline-dependent effects of amphetamine on visual attention, the findings for methylphenidate and atomoxetine suggest important, apparently opposing effects on visual signal detection performance produced via blockade of the DA and NA transporters. The deleterious effects of atomoxetine on performance were especially noteworthy in view of its use as a treatment in ADHD.

## Introduction

Several compounds have been used to treat attentional deficits in attention deficit hyperactivity disorder (ADHD), including psychostimulant drugs such as dextro-amphetamine (AMPH) and methylphenidate (MPH), as well as the non (or atypical) stimulant atomoxetine (ATO) (Spencer et al. 2008; Brown et al. 2009; Epperson et al. 2011; Durell et al. 2013; Frick et al. 2017). AMPH exerts its neurocognitive effects by inhibiting monoamine transporters and driving vesicular dopamine (DA) release, whereas MPH acts as a selective dopamine transporter (DAT) and noradrenaline transporter (NAT) inhibitor, and ATO as a selective NAT transporter inhibitor (Wolraich et al. 2005; Wilens 2006; Bidwell et al. 2011; Del Campo et al. 2011). There are clinical differences in efficacy of ADHD treatments, with some studies reporting superior efficacy of AMPH and occasionally MPH (Del Campo et al. 2011; Durell et al. 2013; Bédard et al. 2015; Bickel et al. 2016; Nagy et al. 2016; Rezaei et al. 2016; De Crescenzo et al. 2017; Cortese et al. 2018; Kowalczyk et al. 2019; Elliott et al. 2020). Supporting these findings, there have been several attempts to investigate the comparative effects of these anti-ADHD drugs on attentional performance and impulsivity in preclinical models with rodents using tasks such as the 5-choice serial reaction time task (5-CSRTT) (Cole and Robbins 1987; Chudasama et al. 2005; Robinson 2012; Caprioli et al. 2015; Caballero-Puntiverio et al. 2017; Higgins et al. 2020), the related 5-choice continuous performance task (5C-CPT) (MacQueen et al. 2018; Young et al. 2020), two-choice visual detection tasks (McGaughy and Sarter 1995; Grilly et al. 1998; Turner and Burne 2016), and a touch-screen continuous performance task (rCPT) (Caballero-Puntiverio et al. 2019, 2020).

When tested in task environments under specific conditions, low-dose AMPH (and to a more limited extent, MPH) improves attentional performance, but impairs performance at higher doses in an inverted U-shaped function, depending on such factors as the baseline level of performance (low levels being enhanced), schedule and rate of stimulus presentations, task employed, and degree of training (Grilly et al. 1998; Sagvolden and Xu 2008; Turner and Burne 2016; Higgins and Silenieks 2017; MacQueen et al. 2018; Caballero-Puntiverio et al. 2020; Higgins et al. 2020; Young et al. 2020). However, beneficial effects of ATO on attentional performance are less evident, with some studies showing improvement with low doses under certain conditions such as lengthened inter-trial intervals or Go/NoGo paradigms (Jentsch et al. 2009; Robinson 2012; Caballero-Puntiverio et al. 2020; Toschi et al. 2021), noradrenergic depletion (Newman et al. 2008), or in young rats (Cain et al. 2011), but others showing impairments (Mar et al. 2017; Higgins et al. 2020; Toschi et al. 2021; Hervig et al. 2023).

Parallel to, and perhaps contributing to, these effects on attention, drugs such as AMPH and MPH generally both dose-dependently and baseline-dependently, increase premature responding (impulsivity) (Carli et al. 1983; Cole and Robbins 1987; Evenden and Ryan 1996; Robbins 2002; van Gaalen et al. 2006a; Robinson et al. 2008; Economidou et al. 2012; Tomlinson et al. 2014; Ding et al. 2018; Toschi et al. 2021; Hervig et al. 2023). These actions are in marked contrast to those of ATO, which generally reduces impulsivity in several test situations (Faraone et al. 2005; Robinson et al. 2008; Jentsch et al. 2009; Paterson et al. 2011, 2012; Baarendse and Vanderschuren 2012; Wilson et al. 2012; Ansquer et al. 2014; Pillidge et al. 2014; Fitzpatrick and Andreasen 2019; Higgins et al. 2020; Toschi et al. 2021).

Recently, the classical theories of signal detection theory (TSD) (McGaughy and Sarter 1995; MacQueen et al. 2018) and visual attention (TVA) (Huang-Pollock et al. 2012; Bundesen et al. 2015; Habekost 2015; Hervig et al. 2023) have been employed to better characterise the effects of stimulant drugs on attentional performance. TSD uses indices of discriminative sensitivity (d’ or A’) as well as perceptual bias (𝛽, B’’) that control for effects of drugs specifically upon attentional function, whilst taking into account their effects on their general tendency to respond. TVA (Bundesen 1990) combines measures of response latency and accuracy to provide more sensitive indices of performance in a theoretical framework for assessing human attentional processes.

Therefore, this study aims to investigate the effect of commonly used ADHD treatments of AMPH, MPH, and ATO on attention while minimising the possible confounding influence of impulsivity utilizing a rapid visual signal detection task (SDT) (Turner et al. 2016). This task enabled analysis using both TSD and TVA approaches, in the former case by measuring correct rejections following non-occurrence of the visual targets, as well as ‘Hits’ upon target presentation. We were also able to vary attentional demand by manipulating the duration of the visual target and to categorise high, medium, and low performing groups to assess baseline-dependent effects. Based on the findings of Turner and Burne (2016), we hypothesised that all three ADHD medications would produce dose-and baseline-dependent effects following their systemic administration. Specifically, we expected low doses to enhance cognitive performance in low-attentive (LA) subjects, while high-attentive (HA) subjects might exhibit dose-independent impairments because of the well-known principle of baseline-or rate-dependent effects that also leads to inverted-U shaped dose-response functions (Cools and Robbins 2004). Furthermore, based on their established clinical efficacy, we anticipated that AMPH would elicit the most pronounced cognitive effects, followed by MPH, with ATO exerting the least beneficial, and possible detrimental, effects.

## Methods

### Compliance with ethical standards

This research has been regulated under the Animals (Scientific Procedures) Act 1986 Amendment Regulations 2012 (Project licence PA9FBFA9F, Neurobehavioural Mechanisms of Mental Health, PPL licence holder: Professor Amy Milton) following ethical review by the University of Cambridge Animal Welfare and Ethical Review Body (AWERB).

### Animals and housing

Twenty-four adult male Sprague Dawley rats (Charles River, Kent, UK) were housed in a room with 21°C ± 2, 60% humidity, and 12h:12h light/dark cycle (lights on at 7pm) and acclimatized for 7 days prior to commencement of procedures. Rats were housed in standard cages with wood-chip bedding and cage enrichment (a cardboard tunnel and woodblock) and kept in groups of four. Food restriction was applied when rats reached ∼300g bodyweight prior to behavioural experiments and restricted to 90% of their free-feeding bodyweight. The amount of food was calculated daily and adjusted to sugar pellet consumption during testing to a total of 18-25g of Purina rodent chow per animal. Water was available *ad libitum* in the home cage, but not during behavioural experiments.

### Behavioural apparatus

Operant chambers (Med Associates, Georgia, VT, USA) were placed in sound-attenuating MDF boxes with ventilation fans, Figure 1. Eight boxes were assembled for the signal detection task, and all operant chambers had the same dimensions (30 x 29 x 24 cm) in sound-attenuating chambers (38 x 60 x 64 cm) equipped with electric fans. The operant chambers for the signal detection task contained a house light, signal light, and nose poke port located centrally of a chamber wall. A food magazine with external pellet dispensers attached were located on either side of the nose poke port and delivered 45 mg sucrose pellets (TestDiet 5TUL, OpCoBe, UK). Both food magazines and the nose poke port were equipped with a light and head entry detector. The software to operate and record data was K-Limbic (Med Associates, St. Albans, VT, USA).

**Figure 1:**
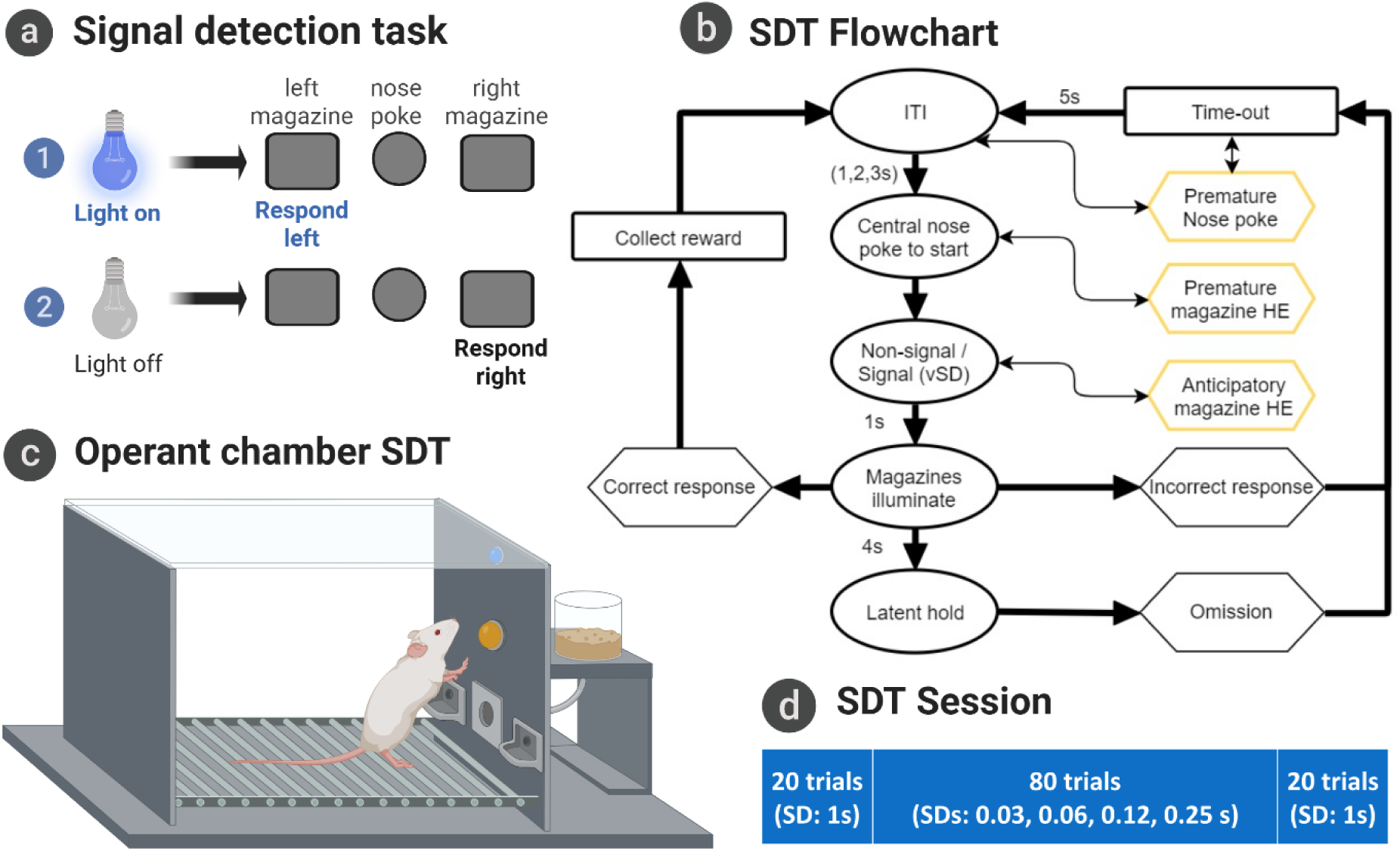
Overview of the signal detection task (SDT). A: Explanation of the SDT; B: A flowchart of SDT trials and outcomes; C: Operant chamber of the SDT; D: Overview of number of trials and variable signal duration in one session of the altered SDT. Abbreviations: HE = Head entry, ITI = Inter Trial Interval, SD = Signal Duration, vSD = variable signal duration. Figure was created in BioRender; Wilod Versprille, L. (2025) https://BioRender.com/8h79hjm.

### Pre-training and Signal Detection task

Pre-training for the SDT has been described in detail by Turner and colleagues (2016) (Figure 1 and Supplementary table S1). Rats received training once or twice daily for 5-7 days a week. In short, rats were first trained to collect sucrose pellets from the two food magazines. Rats were then trained to nose-poke in a central port and subsequently make a head entry in one of the two food magazines to obtain a sugar pellet. The signal detection component of the task was then introduced, where the signal light was either illuminated or extinguished after a central nose poke. The rats were trained to associate one food magazine with the light on (signal trial) and the other with the light off (no signal trial).

Finally, variable signal durations (SDs) were introduced to increase attentional demand. Rats were then trained until they achieved a stable baseline performance. The signal side was counterbalanced and remained constant for each rat. Following the attainment of a stable baseline over 3-5 consecutive days (usually after 10 sessions), pharmacological interventions commenced.

### Drugs

To examine the effects of commonly utilised ADHD medications, subjects received systemic administrations of varying doses of AMPH, MPH and ATO in three subsequent Latin square designs. A drug testing day was preceded by a baseline session and followed by a washout day. Each Latin square was followed by ≥3 additional washout days. Despite studies showing lasting synaptic plasticity effects following ≥1 mg/kg AMPH (Robinson et al. 1982; Kolb et al. 2003; Singer et al. 2009), no such effects have been shown at 0.4 mg/kg, nor did we observe lasting behavioural effects of AMPH. Thus, drug-induced neuroplasticity was unlikely to be a confounding factor in our acute behavioural pharmacology experiments. Drugs were administered systemically via intraperitoneal injection (1 ml/kg) (Terumo® Agani™ 0.5×16 mm needles and Terumo® 1 ml syringes) 30 min prior to testing. Drugs were dissolved in 0.9% sterile saline and injected in various doses, based on published studies. Drugs tested included d-amphetamine hemisulfate salt (0.1; 0.2; 0.4 mg/kg) (Sigma-Aldrich, UK) (Cole and Robbins 1987; Cardinal et al. 2000; van Gaalen et al. 2006b; St Onge and Floresco 2009; Turner and Burne 2016; Turner et al. 2017), methylphenidate hydrochloride (0.3; 1; 3 mg/kg) (Johnson Matthey, UK, GB-D105HL1) (van Gaalen et al. 2006b; Robinson 2012; Fernando et al. 2012; Nishitomi et al. 2018), and atomoxetine hydrochloride (0.1; 0.3; 1 mg/kg) (EDQM, France, Y0001586) (Newman et al. 2008; Robinson 2012; Fernando et al. 2012; Mar et al. 2017; Nishitomi et al. 2018).

### Modelling attention

#### Signal detection theory

TSD provides an analytical framework for assessing performance that separates sensitivity from bias. This method more closely evaluates attention in comparison to the more commonly reported measure of accuracy (Tanner and Swets 1954; Green and Swets 1966; Stanislaw and Todorov 1999). First, the number of Hits, Misses, False Alarms (FAs), and Correct Rejections (CRs) were quantified. A Hit was the correct identification of the presence of the visual cue; a CR reflected a correct response to its absence. Conversely, a Miss occurred when the subject failed to detect a presented cue, and an FA was noted when the subject incorrectly reported a cue in its absence. Then, sensitivity (d’, Equation 1) and response bias (β, Equation 2) were calculated based on signal detection theory, where sensitivity, which more closely reflects attention, reflects the ability to discriminate between signal and noise trials and response bias, a subject’s tendency to report the presence of a signal (i.e. perceptual bias). The parameter β is highly correlated with the other classical TSD parameter for bias C. In contrast to Go/NoGo paradigms, the bias parameter in the SDT is not indicative of a liberal/conservative motor response strategy. It does not only simply quantify left-right lever bias as these were counterbalanced to report signal and no signal (noise). Thus, β measures the tendency to report signal (light on) or noise (light off), i.e. perceptual bias and not general’responsivity’. In the rare cases when a subject had a 𝑝_Hit_or 𝑝_FA_of 0 this would result in a d’ and β of (-)infinite value and so these were converted into missing values to avoid issues with creating averages and statiss.

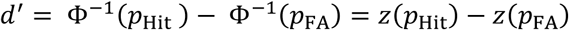

**Equation 1**: Calculation of sensitivity d’, where φ^-1^ = z and returns a z-score. 𝑝_Hit_ and 𝑝_FA_ indicate the probability of making a Hit and False Alarm, respectively. The d’ is in standard deviation units, where 0 indicates a complete inability to distinguish signal from noise, and larger positive values indicate an increasingly greater ability to distinguish signals from noise. Negative values arise through sampling error or response confusion.

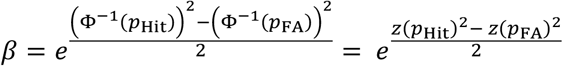

**Equation 2**: Calculation of bias β using the standardized probability of making a Hit and False Alarm. A β value of 1 signifies no bias, β < 1 a bias towards reporting the signal and β > 1 a bias towards reporting its absence.

#### Theory of visual attention

TVA (Bundesen 1990) is another computational model that can be used to analyse and model attentional performance. TVA has a long track record in modelling behavioural data from both neurotypical human populations (Bundesen et al. 2015) and clinical groups (Habekost 2015). More recently, it has also been applied to rodent data from the 5-CSRTT under different pharmacological manipulations (Fitzpatrick et al. 2017, 2019; Hervig et al. 2023). Within the TVA framework, the detection time of a signal is assumed to follow an exponential distributed with a rate parameter 𝑣 (visual processing speed). Under this assumption, the probability of detecting a signal is given by 1 − 𝑒^−𝑣𝜏^, where 𝜏 is the signal duration.

As signal duration increases, so does the probability of detection. Consequently, the probability of not detecting a signal is 𝑒^−𝑣𝜏^. If no signal is detected, we assume that the rat nonetheless makes a signal response with probability 𝑝_s_. A value of 𝑝_s_= 0.5 indicates that the rat is equally likely to make or withhold a signal response when no signal is detected, whereas a value of 𝑝_s_ > 0.5 indicates a bias toward responding. Combining these assumptions, the TVA framework allows us to model the probability of a Hit (𝑝_Hit_), a Miss (1 − 𝑝_Hit_), a False Alarm (𝑝_FA_), and a Correct Rejection (1 − 𝑝_FA_) as follows:

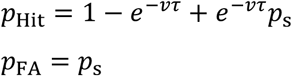

**Equation 3**: TVA-modelling of the probability of making a Hit (𝑝_Hit_), a Miss (1 − 𝑝_Hit_), a False Alarm (𝑝_FA_), and a Correct Rejection (1 − 𝑝_FA_), where 𝑣 is visual processing speed (measured in Hz or signals/second), 𝜏 is the signal duration, and 𝑝_s_ is the probability of making an signal response despite no signal being detected.

Figure 4A presents the model fitted to representative data from a rat performing the SDT with variable SDs. To aid interpretation, the figure also includes a visual representation of the model parameters, illustrating the mathematical framework described above. Parameter estimation was carried out using a custom-build Matlab script and a maximum-likelihood procedure. It is important to note that, unlike previous TVA-based modelling of rodent data from the 5-CSRTT, we do not report the *t*_0_ parameter (preprocessing time) for the SDT. This parameter could not be reliably estimated and was effectively zero in many fits. Consequently, in a second iteration of the modelling, *t*_0_ was fixed at zero.

## Statistical analysis

Drug-induced effects were assessed across multiple behavioural parameters, including response speed, anticipatory responses and accuracy. Additionally, TVA and TSD analyses were conducted to dissociate attentional processes from other factors influencing accuracy, such as perceptual bias. Results were obtained and pre-processed with software provided by K-Limbic (Med Associates) and subsequently exported for data analysis using custom scripts written in RStudio (RStudio Inc., version 2023.06.0, R v4.2.2). Data are presented as accuracy for the overall or subset of trials (the middle block of 80 variable SDs) but also separated by SD. Premature responses are presented as raw values and response latencies in seconds. Results are presented as mean ± S.E.D and significance was set at p < 0.05. Performance groups were created based on low, middle and high baseline performing groups, each comprising a third of the ranked accuracy scores averaged across three baseline sessions (Figure 2A) or the four baseline sessions within each Latin square design. Baseline performance groups remained stable, with only small changes observed where subjects moved from low to middle or middle to high, but never from low to high baseline performance groups. Prior to statistical analysis the assumptions, e.g. normality, were checked. When normality was violated, either non-parametric tests were used or data were transformed. Spearman correlations were used to investigate the strength of the relationship between variables. To compare the effect of the drug dosing groups on the behavioural outcomes and mathematical models, mixed-effects models (with Satterthwaite’s method) of varying magnitudes were performed. When a mixed-effects model was run, the restricted maximum likelihood (REML) was used for statistical models that included individual performance groups as a random effect, when performance subgroups were included as a fixed effect the maximum likelihood estimation was used instead. The mixed-effects models could include the following independent variables: performance group (between-subjects) as fixed or random effect, drug dose (within-subject), signal duration trials (within-subject) and rat ID as random effect to accommodate for repeated measures, the F and p-values of the main dose effect and dose x performance level interaction can be found in Table 2. Pairwise *post-hoc* comparisons were performed where appropriate, using False Discovery Rate (FDR) for p-value adjustment. ∗p < 0.05, **p < 0.01, ***p < 0.001, ****p < 0.0001. Details of the outcome of *post-hoc* analyses are provided in the Supplementary Material.

**Figure 2:**
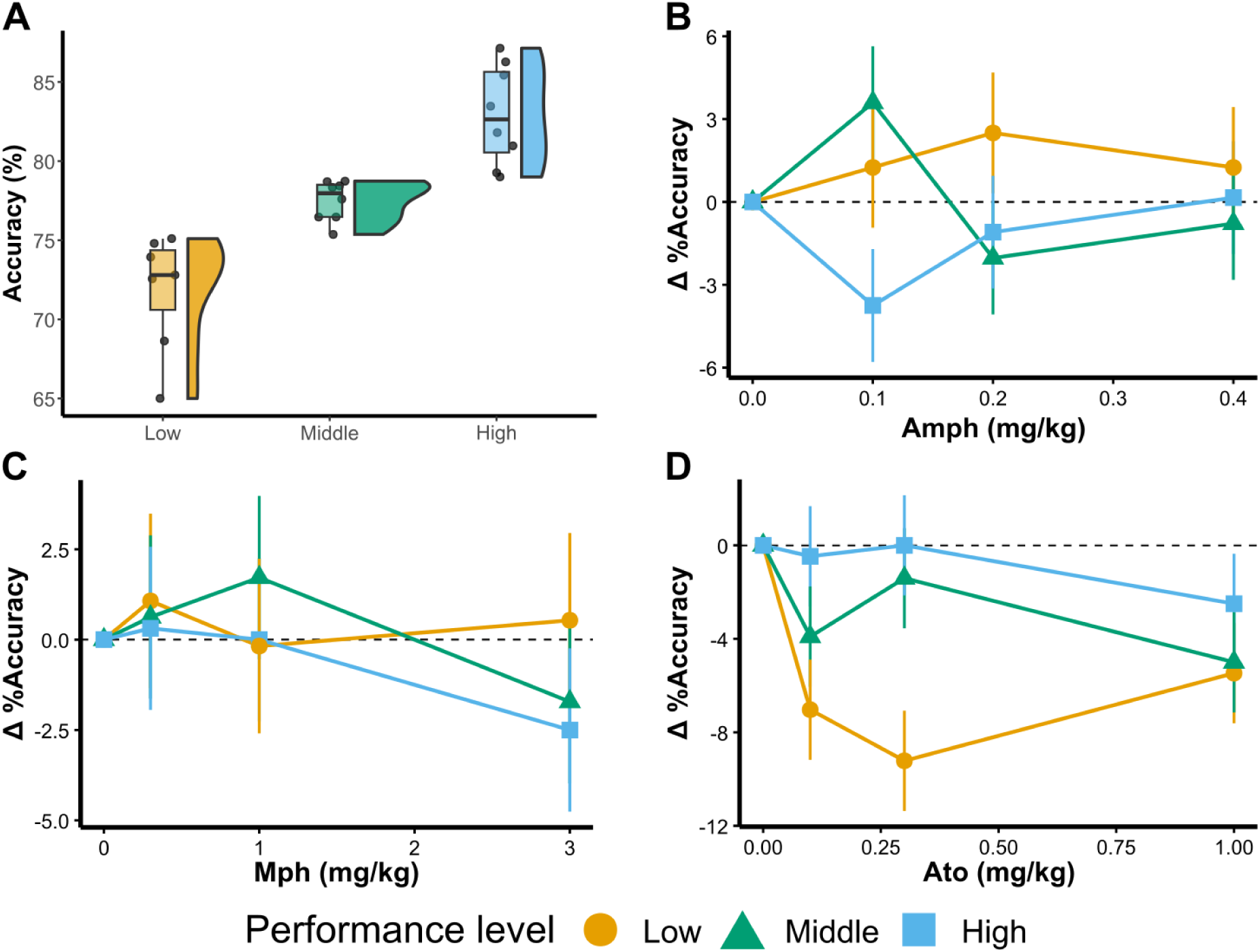
Baseline and dose-dependent effects of systemic anti-ADHD drug administration on the SDT. Distribution of performance levels (n = 23) of average baseline performance in the 3 sessions prior to drug testing (A). The drug effects on accuracy of the middle 80 trials depicted as the mean ± SED (standard error of the differences of the means) between vehicle and drug for each performance groups for AMPH, n = 23 (B), MPH, n = 23 (C) and ATO, n = 24 (D).

**Table 1:** The Spearman correlation coefficients between the behavioural and mathematical parameters of the SDT (n = 24). Abbreviations and variable names: d’ = sensitivity; β = Beta, bias; *v* = visual processing speed; *p*s = probability of making a signal response despite no signal being detected; *p*Hit = probability of making a hit; *p*FA = probability of making a false alarm.

| Mean Baseline | $d'$ | | $\beta$ | | $v$ | | $p_s$(guessing) | |
| --- | --- | --- | --- | --- | --- | --- | --- | --- |
|  | R-value | p-value | R-value | p-value | R-value | p-value | R-value | p-value |
| Accuracy | 0.92 | <0.001 *** | -0.48 | 0.02 * | 0.86 | <0.001 *** | -0.37 | 0.07 |
| $d'$ | | | -0.2 | 0.35 | 0.74 | <0.001 *** | -0.67 | <0.001 *** |
| $\beta$ | -0.2 | 0.17 | | | -0.61 | 0.001 ** | -0.43 | 0.04 * |
| $v$ | 0.74 | <0.001 *** | -0.61 | 0.001 ** | | | -0.25 | 0.25 |
| $p_s$ (guessing) | -0.67 | <0.001 *** | -0.43 | 0.04 * | -0.25 | 0.25 | | |
| $p_{Hit}$ | 0.47 | 0.02 * | -0.87 | <0.001 *** | 0.84 | <0.001 *** | 0.20 | 0.35 |
| $p_{FA}$ | -0.81 | <0.001 *** | -0.31 | 0.14 | -0.32 | 0.13 | 0.93 | <0.001 *** |
| Trial initiation latency | -0.22 | 0.29 | 0.47 | 0.02 * | -0.37 | 0.08 | -0.09 | 0.69 |
| Correct Response latency | -0.04 | 0.85 | 0.12 | 0.58 | -0.21 | 0.33 | -0.17 | 0.44 |
| Incorrect Response Latency | 0.04 | 0.86 | 0.09 | 0.69 | -0.07 | 0.76 | -0.23 | 0.27 |
| Anticipatory responses | -0.1 | 0.64 | -0.25 | 0.24 | 0.08 | 0.73 | 0.06 | 0.77 |

**Table 2:** The statistical results from d-amphetamine, methylphenidate and atomoxetine administrations. The following linear mixed effects model was applied to obtain the main dose effect and dose x performance interaction: Variable ∼ dose * performance level + +(1|ratID).

|  | Amphetamine |  |  |  | Methylphenidate |  |  |  | Atomoxetine |  |  |  |
| --- | --- | --- | --- | --- | --- | --- | --- | --- | --- | --- | --- | --- |
|  | Dose<br>(Df 3, 69) |  | Dose x<br>performance<br>(Df 6, 69) |  | Dose<br>(Df 3, 69) |  | Dose x<br>performance<br>(Df 6, 69) |  | Dose<br>(Df 3, 72) |  | Dose x<br>performance<br>(Df 6, 72) |  |
|  | F | P | F | P | F | P | F | P | F | P | F | P |
| **Accuracy** | 0.10 | n.s. | 2.69 | 0.02 | 0.95 | n.s. | 0.39 | n.s. | 5.80 | 0.001 | 2.60 | 0.02 |
| **$d'$** | 0.49 | n.s. | 2.87 | 0.01 | 1.93 | n.s. | 0.53 | n.s. | 2.89 | 0.04 | 1.86 | 0.10 |
| **$\beta$** | 3.78 | 0.01 | 1.84 | 0.1 | 2.18 | 0.10 | 0.64 | n.s. | 1.32 | n.s. | 0.50 | n.s. |
| **$v$** | 0.53 | n.s. | 1.61 | n.s. | 2.07 | n.s. | 0.50 | n.s. | 4.09 | 0.01 | 1.96 | 0.08 |
| **$p_s$** | 0.83 | 0.48 | 2.59 | 0.03 | 3.75 | 0.01 | 1.57 | n.s. | 0.56 | n.s. | 1.05 | n.s. |
| **No signal** | 1.16 | n.s. | 1.44 | n.s. | 4.29 | 0.008 | 0.36 | n.s. | 0.09 | n.s. | 1.26 | n.s. |
| **30 ms** | 1.22 | n.s. | 1.09 | n.s. | 2.43 | 0.08 | 1.15 | n.s. | 3.13 | 0.03 | 0.83 | n.s. |
| **60 ms** | 1.06 | n.s. | 0.85 | n.s. | 1.48 | n.s. | 0.39 | n.s. | 1.85 | n.s. | 1.91 | 0.09 |
| **120 ms** | 0.31 | n.s. | 0.54 | n.s. | 0.57 | n.s. | 0.48 | n.s. | 1.40 | n.s. | 1.98 | 0.08 |
| **250 ms** | 0.32 | n.s. | 0.38 | n.s. | 1.72 | n.s. | 2.03 | 0.07 | 0.07 | n.s. | 1.22 | n.s. |
| **1 s** | 1.51 | n.s. | 2.17 | 0.06 | 0.78 | n.s. | 0.94 | n.s. | 0.24 | n.s. | 0.53 | n.s. |
| **Trial initiation latency** | 6.70 | <0.001 | 0.30 | n.s. | 2.23 | 0.09 | 2.41 | 0.04 | 1.41 | n.s. | 0.53 | n.s. |
| **Response latency** | 0.76 | n.s. | 1.02 | n.s. | 0.38 | n.s. | 0.80 | n.s. | 1.09 | n.s. | 0.74 | n.s. |
| **Anticipatory responses** | 4.36 | 0.007 | 0.52 | n.s. | 2.95 | 0.04 | 0.38 | n.s. | 9.67 | <0.001 | 0.52 | n.s. |

## Results

### Validation of mathematical modelling of attentional processing and perceptual bias

To dissociate perceptual and attentional mechanisms, two mathematical models were applied: TSD quantified sensitivity and decision (perceptual) bias, while TVA estimated attentional capacity and processing speed. Unlike TSD, which measures signal–noise discrimination, TVA provides a mechanistic account of how attention is allocated and how prone subjects are to guess. We assessed the overlap between these parameters, as well as their relationship with accuracy, using correlational analyses (Table 1). The TVA and TSD effectively captured underlying attentional processes, as evidenced by significant positive correlations between accuracy, d’ and *v*, while these measures showed no significant relationship with response speed, trial initiation speed, or anticipatory responses. In line with expectations (see Equations 1&3), d’ was significantly positively correlated with p_Hit_, and negatively with p_FA_, whereas visual processing speed (*v*) was only significantly correlated with p_Hit_.

Furthermore, parameters modelling perceptual bias (β) and guessing (*p*_s_) were significantly negatively correlated, indicating that subjects with a stronger bias to reporting signal exhibited a greater tendency to guess. Moreover, d’ but not *v,* was negatively correlated with guessing, while β was correlated with accuracy and *v* but not d’. This is likely due to the strong correlation of d’ and guessing with the false alarm rate (p_FA_), and on the other hand strong correlation for β, accuracy, and *v* with the hit rate (p_Hit_), as expected (see Equations 1-3). Additionally, β was positively correlated with trial initiation latency, though not the number of anticipatory responses, both parameters that are commonly associated with motivation. Hence, the more conservative tendency (i.e. to report “no signal”) is associated with a slower tendency to initiate the trial. In contrast to the negative correlation presented in Table 1, there appeared to be a positive relationship between β and accuracy in Figure 3B. While the negative correlation slightly decreased over time (average AMPH baseline performance r =-0.21, p = 0.04) and eventually disappeared (average ATO baseline performance r = 0.02, p = non-significant), the positive relationship observed in Figure 3B likely resulted from an anomalous vehicle effect.

**Figure 3:**
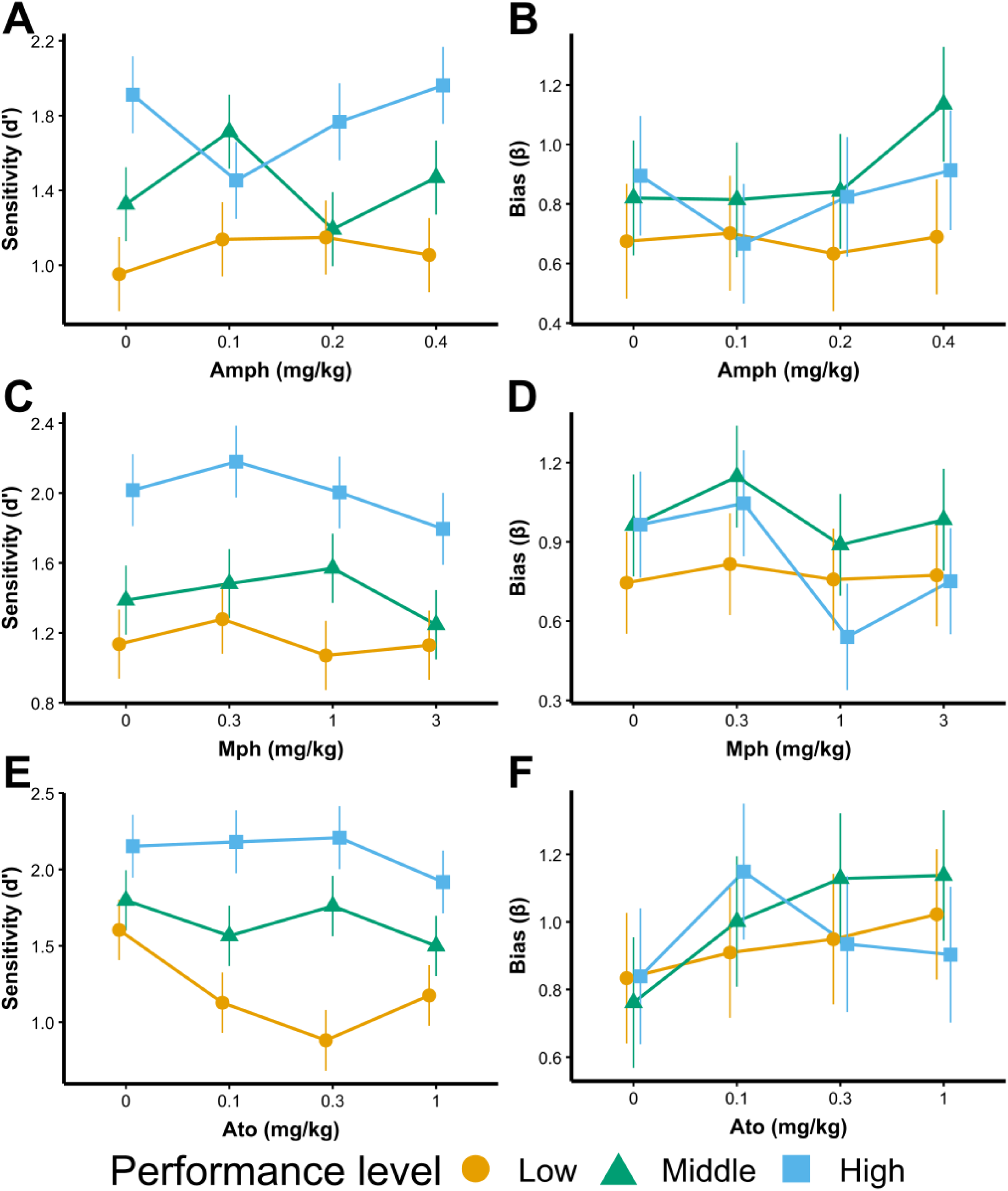
Baseline and dose-dependent effects of systemic anti-ADHD drug administration on the TSD parameters discriminability and perceptual bias (mean ± SED) in each performance group. Baseline and dose-dependent effects of systemic AMPH (n = 23) on discriminability (A) and perceptual bias (B), systemic MPH (n = 23) on discriminability (C) and perceptual bias (D), and systemic ATO (n = 24) on discriminability (E) and perceptual bias (F).

**Figure 4:**
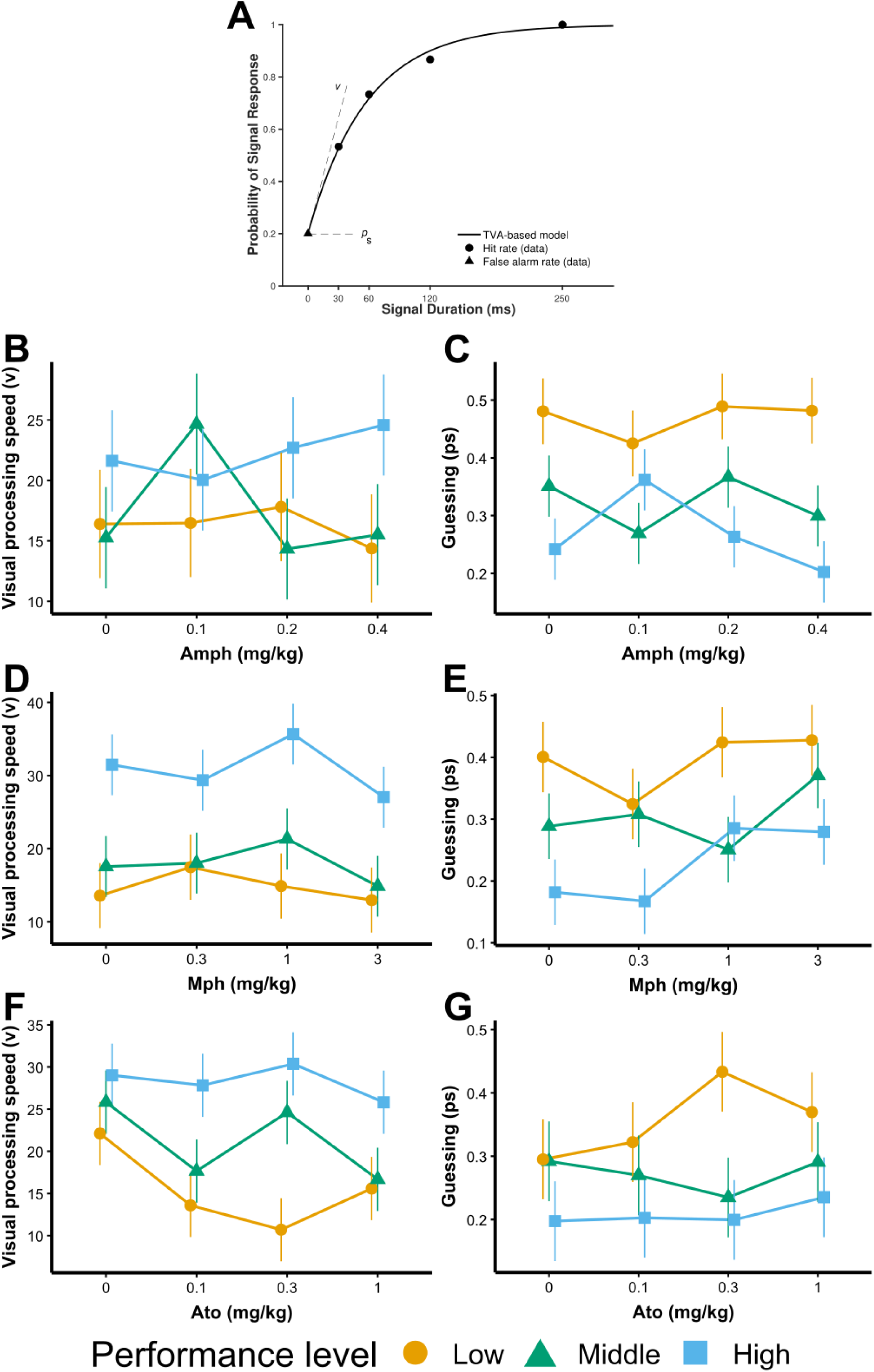
Baseline and dose-dependent effects of systemic anti-ADHD drug administration on the TVA parameters visual processing speed (v) and guessing (*p*s) (mean ± SED) in each performance group. (A) TVA model fitted to representative data from a single SDT session with variable signal duration. The observed false alarm rate (triangle) and hit rates (dots) are shown alongside the model’s predicted probability of a signal response (solid line) as a function of the signal duration. The figure also displays the model parameters: *p*s represents the probability of making a signal response despite no signal being detected, illustrated by the horizontal line indicating the false alarm rate when no signal is presented (i.e., a signal duration of zero ms). v denoted the visual processing speed of signal detection, illustrated as the initial slope of the curve at zero ms signal duration. AMPH dose-and baseline-dependent (n = 23) modulation of visual processing speed (B) and the willingness to guess (C). MPH dose-and baseline-dependent modulation (n = 23) of visual processing speed (D) and guessing (E). ATO dose-and baseline-dependent modulation (n = 24) of visual processing speed (F) and willingness to guess (G).

In summary, these results suggest that TVA and TSD model complementary, albeit partially overlapping, processes. This led to, for example, the finding that more guessing was related to poorer discriminability. In further support of their complementarity was the differing way anti-ADHD drugs affected these processes (see below). Of particular interest is the difference between low and high baseline performing subjects (Figure2A). The correlations were consistent with the strongest differentiation between low and high baseline performing subjects being changes in d’ and v, but not β or guessing. This indicates that the difference was rooted in differential attentional capabilities of these sub-groups, and not perceptual bias or changes in motor response strategy. Therefore, the baseline performance groups will be referred to as low attentive (LA), middle attentive (MA) and high attentive (HA).

### Low-dose amphetamine modulated attention in a baseline-dependent manner

Systemic administration of low-dose AMPH (0.1, 0.2, 0.4 mg/kg) produced baseline-dependent effects (Table 2), enhancing accuracy (Figure 2B) and d’ (Figure 3A), but not visual processing speed (Figure 4B), in LA and MA subjects, while impairing HA subjects. These effects appeared to be driven by the 0.1 mg/kg dose, although the AMPH effects did not quite survive rigorous *post-hoc* testing (Supplementary table S2). Furthermore, AMPH appeared to selectively enhance LA accuracy for 1s signal trials (trend for dose x baseline performance interaction, Table 2), baseline-dependently modulate no signal (LA improved) and ambiguous shorter SDs (LA impaired) trials (Supplementary table S2).

AMPH dose-dependently increased perceptual bias towards reporting no signal (Figure 3B; β dose effect; F_(3, 69)_ = 3.78, p = 0.01), driven by an increase in conservative perceptual bias at 0.4 mg/kg. Although this appears specific for MA and HA (Figure 3B), only a trend towards a baseline-dependent effect on perceptual bias was observed from the dose x baseline performance interaction term (Table 2). On the other hand, there was a significant baseline-dependent modulation of the probability of reporting signal in the absence of a stimulus (i.e., the willingness to guess). This appeared to be driven by a decreased willingness to guess at 0.1 mg/kg for LA subjects but an increase for HA subjects (Figure 4C). Nonetheless, the AMPH effects on perceptual bias and willingness to guess also did not quite survive rigorous *post-hoc* testing. AMPH dose-dependently induced a motor speeding effect for all subjects, irrespective of baseline performance and was characterized by significantly faster trial initiation latency (dose effect F_(3, 69)_ = 6.70, p < 0.001) and an increased number of anticipatory responses at the food magazines (dose effect F_(3, 69)_ = 4.36, p = 0.007). Response latency on the other hand was unaffected.

### Effects of methylphenidate on signal detection task

MPH did not exhibit dose-or baseline-dependent effects on accuracy, although high-dose MPH appeared to impair performance in HA subjects (Figure 2C, Table 2, and Supplementary Table S3). Consistent with these findings, no significant effects were observed for sensitivity (d’) or visual processing speed, although perceptual bias (β) showed a trend to decrease (becoming more liberal and hence likely to report the presence of a signal) (Table 2. Figure 3C-D and Figure 4D). However, when assessing performance across the entire SDT session rather than specifically the vSD block of 80 trials, a trend toward impaired performance following high-dose MPH emerged (dose effect: F_(3, 69)_ = 2.44, p = 0.07). No significant interactions with SDs were found, although high-dose MPH appeared to enhance accuracy on ambiguous 30 ms SD, while impairing accuracy on other SDs and in the no signal trials (see Supplementary Table S3). MPH administration significantly increased guessing (Figure 4E; *p*_s_ dose effect; F_(3, 69)_ = 3.75, p = 0.01), particularly at higher doses, suggesting a greater likelihood of responding before adequate stimulus processing. Additionally, MPH significantly increased anticipatory responses (F_(3, 69)_ = 2. 95, p = 0.04) and tended to speed trial initiation (dose effect F_(3, 69)_ = 2.23, p = 0.09), particularly for LA and MA subjects (dose x baseline performance F_(6, 69)_ = 2.41, p = 0.04), although these effects did not quite survive rigorous *post-hoc* testing. Response latency remained unaffected by MPH.

### Atomoxetine impaired attention in a dose-and baseline-dependent manner

Unlike AMPH and MPH, ATO significantly impaired accuracy in a dose-dependent (dose effect F_(3, 72)_ = 5.80, p = 0.001) and baseline-dependent manner (dose x baseline performance F_(6, 72)_ = 2.6, p = 0.02). Specifically, low-dose ATO selectively impaired LA subjects, whereas high-dose ATO impaired accuracy in the entire sample (Figure 2D, Table 2, and Supplementary table S4). This reduction in accuracy was driven by a significant decrease in d’ (Figure 3E; dose effect F_(3, 72)_ = 2.89, p = 0.04) and visual processing speed (Figure 4F; dose effect F_(3, 72)_ = 4.09, p = 0.01), with a trend towards a dose x baseline performance interaction for both d’ (F_(6, 72)_ = 1.86, p = 0.10) and visual processing speed (F_(6, 72)_ = 1.96, p = 0.08). ATO appeared to exert a greater impact on more ambiguous shorter SDs and no signal trials. Despite these impairments in attentional processing, ATO did not significantly alter perceptual bias (β; Figure 3F), the tendency to guess (*p*_s_; Figure 4G), trial initiation latency or response latency (Table 2). However, it did reduce the number of anticipatory responses (dose effect F_(3, 72)_ = 9.67, p < 0.001).

## Discussion

This study utilised the SDT to investigate the dose-and baseline-dependent effects of commonly used ADHD treatments on sustained visual attention in rodents. We found predicted baseline-dependent effects of AMPH on SDT performance, whereby poorer (LA and MA) performers were improved and high performers were impaired, albeit at a lower dose (0.1mg/kg) than in the earlier studies of Turner and colleagues (Turner and Burne 2016; Turner et al. 2017). However, while MPH surprisingly did not significantly alter many aspects of SDT performance, ATO importantly significantly impaired SDT performance by reducing attentional accuracy and sensitivity measures. The LA subjects were significantly impaired at lower doses of ATO, in direct contrast with the enhancing effects of low dose AMPH. Moreover, the MA and HA-performance groups also were mainly impaired, though by higher doses of ATO. All drugs had expected effects on motor responses: AMPH and, (to a more limited extent) MPH, dose-dependently speeded trial initiation latency and increased premature, anticipatory responses without affecting choice response latency. By contrast ATO reduced anticipatory responses, without affecting the latency measures.

### The TSD and TVA provided complementary perspectives on visual attention and decision-making

Two complementary mathematical models were employed to delineate attentional and perceptual mechanisms. The TDS quantifies perceptual sensitivity and perceptual bias, where sensitivity approximates attentional performance without characterising underlying attentional mechanisms. The TVA on the other hand, provides a mathematical framework for understanding how attentional resources are allocated to visual stimuli based on processing speed and the willingness to guess. While TSD evaluates how well subjects distinguish signal from noise (no signal trials), TVA offers insight into the capacity and efficiency of attentional processing. Whereas *v* and d’ captured similar components of accuracy (likely via the hit rate), they could each be influenced independently (via differential involvement of the false alarm rate). Moreover, β and *p*_s_ represented distinct perceptual biases, as their respective dependencies included the hit rate and false alarm rate.

The attentional effects produced by the ADHD treatments were not always uniformly modulated by mathematical parameters of the TSD and TVA modelling frameworks and other behavioural measures. ATO both retarded visual processing speed and reduced discriminative sensitivity, but did not affect responsivity, perceptual bias or the tendency to guess. By contrast, the baseline-dependent effects of AMPH on sensitivity were mirrored by its effect on guessing, but were not accompanied by altered visual processing speed, and perceptual bias was only affected at the higher dose. While not affecting sensitivity or visual processing speed, MPH did significantly increase guessing with a tendency for a more liberal perceptual bias. The effects on guessing may correspond to some numerical evidence of impaired accuracy and discriminative sensitivity at the highest dose of MPH (Figures 2C, 3C), suggesting that guessing may represent a more sensitive index of impairment in certain situations. This substantiates the notion that the TVA and TSD provide complementary perspectives on how drugs may affect visual attention and perceptual decision-making.

### Effects of stimulant drugs AMPH and MPH on attentional performance

The improvement in SDT performance produced by AMPH in low performing rats is consistent with pro-cognitive effects of this drug reported both clinically and preclinically (Puumala et al. 1996; Bizarro et al. 2004; Robinson 2012; Fitzpatrick et al. 2017; Caballero-Puntiverio et al. 2017, 2019, 2020; Nishitomi et al. 2018; Higgins et al. 2020; Toschi et al. 2021). However, findings have been inconsistent, with some studies failing to observe overall or baseline-dependent cognitive enhancement, and others reporting (often baseline-dependent) cognitive impairments (Jentsch et al. 2009; Koffarnus and Katz 2011; Navarra et al. 2017; Caballero-Puntiverio et al. 2020; Hervig et al. 2023). Noteworthy is the observation that dose and baseline-dependent cognitive modulation of AMPH with two-choice signal discrimination tasks (c.f. also Turner and Burne 2016) have produced more consistent accuracy enhancing effects than with the 5-CSRTT (see for example, Cole and Robbins 1987). However, the enhanced sensitivity parameter following low dose AMPH in the SDT is also consistent with its enhancement of 5C-CPT and rCPT performance in rats, mice and humans (MacQueen et al. 2018; Caballero-Puntiverio et al. 2019, 2020; Young et al. 2020). The presence of no signal trials in both the present SDT and in the 5C-CPT may explain their greater sensitivity to AMPH-induced attentional improvement than in the 5-CSRTT.

Attention-enhancing effects of MPH in previous studies, for example, in the 5-CSRTT and 5C-CPT, have generally been marginal, and generally in low performing animals only (e.g. Paterson et al 2011; Robinson 2012; Caballero-Puntiverio. et al 2017; Navarra. et al 2017). We failed to find beneficial effects of MPH on sensitivity or visual processing speed even in the LA rats and over a reasonable wide dose range, although there was some evidence of enhanced accuracy for no signal and ambiguous short SD trials (e.g. 30 ms). Hervig and colleagues (2023) also found no effect of MPH on accuracy in the 5-CSRTT but did observe enhanced visual processing speed. Furthermore, Hervig and colleagues (2023) reported an increased tendency to guess under uncertainty by MPH, as also found in the present study. Overall, the present behavioural findings are consistent AMPH being more potent than MPH in its neurochemical actions (Kuczenski and Segal 1997).

### Detrimental effects of ATO on attentional performance

In contrast with the stimulants, the non-stimulant drug ATO impaired SDT performance and reduced the number of anticipatory responses. The decreased accuracy was characterised by reductions in discriminative sensitivity and slowed visual attentional processing (the latter finding being consistent with that of Hervig and colleagues (2023) in the 5-CSRTT). The lack of a general slowing of response latencies, or effects on guessing or perceptual bias, suggest a rather specific detrimental effect by ATO on visual attentional function.

Overall, these detrimental effects of ATO contrasted not only with those of the stimulant drugs, but also with several studies reporting no effect (Blondeau and Dellu-Hagedorn 2007; Robinson et al. 2008; Koffarnus and Katz 2011; Robinson 2012; Ansquer et al. 2014; Redding et al. 2019) or even enhanced attentional performance by ATO (Jentsch et al. 2009; Tomlinson et al. 2014; Nishitomi et al. 2018; Caballero-Puntiverio et al. 2019, 2020; Higgins et al. 2020). Those studies reporting enhanced effects of ATO on attention in healthy rodents are generally under specific conditions of low baseline attentional performance (Tomlinson et al. 2014), dependent on specific task parameters such as lengthened inter-trial intervals (e.g. Jentsch et al. 2009), a Go/NoGo paradigm (Caballero-Puntiverio et al. 2020) or the inference of reduced attentional’lapses’ based on less skewed reaction time distributions (Redding et al. 2019). In the latter study, this effect was not modulated by distracting task conditions. In the 5C-CPT task used by Tomlinson and colleagues (2014) the high performing attentional subgroup was impaired by high dose ATO, consistent with the present findings. Additionally, they found that a high impulsivity subgroup showed reductions in premature responses and False Alarms in association with improvements in attentional accuracy. Apparently less concordant with the present results, they found that 2 mg/kg ATO improved sensitivity in low attentive female rats, but no data were shown to indicate whether the drug in parallel reduced the high baseline premature responses and false alarm rates of LA rats in this Go/NoGo test procedure, rather than increasing signal detection (Hits).

This effect of atomoxetine to reduce impulsivity, as measured by premature responses on the 5-CSRTT (and probably by similarly anticipatory responses in the present SDT), has previously been shown in an overwhelming number of studies (e.g. Faraone et al. 2005; Blondeau and Dellu-Hagedorn 2007; Robinson et al. 2008; Paterson et al. 2011, 2012; Robinson 2012; Baarendse and Vanderschuren 2012; Ansquer et al. 2014; Fitzpatrick and Andreasen 2019; Higgins et al. 2020; Toschi et al. 2021). Such effects, especially at longer inter-trial intervals, are sometimes associated with improved accuracy (Jentsch et al. 2009; Robinson 2012; Wilson et al. 2012). However, at shorter inter-trial intervals, where there is less requirement for behavioural inhibition, the opposite effect of impaired accuracy is frequently observed (Jentsch et al. 2009; Higgins et al. 2020).

The effects of ATO to reduce impulsivity could thus also serve to explain the reported increased accuracy (i.e. reduced False Alarms) on NoGo trials in a Go/NoGo stimulus detection task (Bari et al. 2009; Higgins et al. 2020) and in continuous performance tasks (Tomlinson et al. 2014; Caballero-Puntiverio et al. 2019). The present two-alternative forced choice task, however, minimises the need to exert behavioural inhibition and crucially requires an active response to register a correct rejection: this is the likely critical requirement that explains the present ATO deficit, compared with the apparent improvements or no effects seen in other studies. There are also other notable differences between the SDT and tasks like the 5-CSRTT, which involve demands on divided spatial attention rather than vigilance.

Several other studies report similar ATO-induced impairment, for example, on the rodent continuous performance (Mar et al. 2017) and the attentional set-shifting tasks (Newman et al. 2008). In addition to the general dose-dependent impairment by high-dose ATO, LA subjects were affected more and at lower ATO doses than HA subjects, possibly arising from differential neurochemical baselines, in line with baseline-dependent modulation of catecholamines and its metabolites in the striatum and prefrontal cortex (Newman et al. 2008; Rose et al. 2010; Cain et al. 2011; Turner et al. 2017; Broadway et al. 2018; Bensmann et al. 2020; Bayram et al. 2020). For example, it could be speculated that LA subjects may have high levels of tonic NA activity at baseline, which has been associated with poor attentional performance (Aston-Jones et al. 2000) and ATO treatment enhances this tonic state still further.

The marked detrimental effect of ATO on LA (but otherwise intact) rats apparently contrasts with apparent examples of improvements in attentional performance in populations of rodents subjected to prenatal stress (Wilson et al. 2012), NK-1 receptor knockout (Pillidge et al. 2014), noradrenergic depletion (Newman et al. 2008) or in young animals (Cain et al. 2011). Therefore, it is possible that ATO may remediate attention in some pathological populations, although their standing as models of ADHD is more debatable. Perhaps even these improvements depend in part on reductions in impulsive responding.

### Implications of contrasting behavioural effects of MPH, AMPH and ATO for underlying neurochemical mechanisms

The contrasting effects of MPH versus ATO have not previously been reported in a selective and sustained attentional task. The discrepancy could correspond to the lower clinical efficacy reported by some studies for ATO as compared with MPH (Bédard et al. 2015; Kowalczyk et al. 2019), but not by others (Rezaei et al. 2016). Alternatively, the discrepancy could reflect a difference between stimulant and non-stimulant compounds, as ATO reduces anticipatory responses and impulsivity, whilst MPH is associated with increased impulsivity and premature responses (Robinson et al. 2008; Economidou et al. 2012; Tomlinson et al. 2014; Ding et al. 2018). However, these hypotheses do not explain the increased sensitivity to ATO observed for the LA subjects.

Despite our understanding of the pharmacodynamics of these compounds, the precise mechanism and brain regions through which they enhance cognitive functioning are still poorly understood. MPH (as well as AMPH) could influence attention through both DA and noradrenaline (NA) signalling (Berridge et al. 2006). The cognitive enhancing effects of MPH and AMPH have frequently been associated with their dopaminergic effects, although some studies suggest that there is also a noradrenergic influence (Navarra et al. 2017). The dissociation of effects of ATO and MPH could also be, at least partially, mediated by the contrasting functions of cortical and subcortical DA, as catecholaminergic systems converge in the prefrontal cortex (PFC), yet diverge in the striatum (Madras et al. 2005). Due to the great variability in DAT and NAT expression patterns, with high levels of NAT expression in the medial PFC and high levels of DAT expression in the striatum, NAT is primarily responsible for DA reuptake in the medial PFC (Bymaster et al. 2002). Thus, both MPH and ATO raise extracellular catecholamine levels within the PFC, although MPH also increases DA levels in the striatum, while ATO does not (Kuczenski and Segal 1997, 1999; Volkow et al. 2001; Gilbert et al. 2006). Therefore, systemic ATO affects both general NA and cortical DA levels, while MPH affects subcortical DA more substantially (Bymaster et al. 2002). This therefore implies that the differences between MPH and ATO arise because MPH increases subcortical DA activity. Cognitive modulation following changes in NA levels are likely mediated through α1-and α2-adrenoreceptors, which have been shown to affect attentional control (Arnsten 2009; Hervig et al. 2023), while an increase in DA levels likely affects D1 and D2-like receptors. Indeed, Hervig and colleagues (2023) suggest that ATO primarily exerts its effects through the activation of α1-adrenoreceptors.

The differential effects of MPH and ATO could be mediated through opponent DA-NA processes; the profile of effects produced by MPH and AMPH, such as increased impulsivity, (as well as any effects to improve attention) could be ascribed to their pro-dopaminergic actions, whereas the contrasting profile of effects produced by ATO, i.e. reduced impulsivity and impaired attention could be attributed to its pro-noradrenergic actions. In line with this hypothesis, DA-NA interactions would thus be complementary as DA transients are involved in increasing signal, whilst NA transients tend to reduce noise (Arnsten 2000). The implication for understanding the actions of AMPH and MPH is that their parallel actions on the noradrenergic pathways regulate effects of DA mechanisms. A qualitative difference in the roles of DA and NA for behavioural control processes was suggested by Cole and Robbins (1987), as cortical NA depletion following a dorsal noradrenergic bundle lesion neither affected accuracy nor impulsivity, although it did impair response accuracy following systemic AMPH administration (Cole and Robbins 1987; Economidou et al. 2012; Liu et al. 2015). Thus, NA appeared to protect attention from deficits produced by high doses of AMPH presumably caused by excess striatal DA transmission. Moreover, in the present study, a complementary action appears to occur when comparing the overall lack of effects of MPH, (which affects both DAT and NAT), in the LA group with the detrimental effects of ATO (affecting the NAT); the effect on DAT following MPH may protect this subgroup from the deficits produced by ATO. Finally, although the mathematical methods of analysis provided additional parameters for understanding the effects of these compounds, such as visual processing speed and guessing, their utility for identifying underlying neurochemical mechanisms remains to be explored.

ATO may have effects to reduce motivation, based on detrimental effects on breakpoint on progressive ratio schedules produced by doses higher than 0.5 mg/kg, (Higgins et al. 2020) perhaps because of inhibitory actions via action on NAT on mesolimbic DA function - which may also account for its anti-impulsivity action. However, the present detrimental effects on attentional performance in the LA subgroup occurred at lower doses of ATO, specifically affecting d’, and so appear unlikely to be solely motivational in nature.

## Limitations

A frequent concern in interpreting baseline-dependent behavioural effects is regression to the mean. To reduce this possibility, appropriate repeated-measures statistical tests were used to account for within-subject variability, baseline-performance was calculated across baseline tests to differentiate LA from HA subjects, and drug effects were compared to a vehicle treatment (Robbins and Sahakian 1979; Barnett et al. 2005). Our findings warrant further investigation into the potentially clinically detrimental effects of ATO at all doses. We should, however, acknowledge that this study only investigated acute dosing effects, whereas chronic dosing is used in the treatment of ADHD.

## Conclusion

This study replicated and extended the dose-and baseline-dependent cognitive enhancing effects of systemic AMPH, as reported previously on SDT performance (Turner and Burne 2016) and provided further evidence of impaired attention following acute ATO that contrasted with the effects of both AMPH and MPH. The findings suggest that the therapeutic efficacy of ATO may arise primarily from reductions in impulsivity rather than enhancements in attentional processing. Additionally, this study demonstrates that applying frameworks like TSD and TVA - traditionally reserved for clinical assessment - can enhance mechanistic understanding of pharmacological interventions in preclinical models (Habekost 2015). The findings emphasize divergent neuromodulatory effects on behavioural performance of currently available ADHD medications.

## Acknowledgements

We acknowledge Shionogi & Co., Ltd for funding the research. We are grateful to Dr. Karly Turner for her expertise in establishing the SDT task in Cambridge and to Professor Amy Milton as the Home Office licence holder for the regulated procedures used in this research.

## Authors Contributions

LJFWV designed and conducted experiments, collected data, wrote data analysis scripts, analysed data and wrote the manuscript. KY designed experiments, set up equipment and wrote the manuscript. AP wrote mathematical data analysis scripts and wrote the manuscript. JWD designed experiments and wrote the manuscript. TWR secured funding, designed experiments and wrote the manuscript.

