## Supplementary data for "Detrimental effects of atomoxetine on visual signal detection in rats: Comparison with ADHD psychomotor stimulant drugs"

**Table S1:** Overview of the pre-training stages of the signal detection task (SDT). *The 1 second trials in this stage only occurs 10x during the first 20 trials and 10x during the final 20 trials, in the middle segment of 80 trials the other variable signal durations are applied.

| ***Stage*** | ***Level*** | ***Trials (#)*** | ***Session (min)*** | ***Stim Dur (s)*** | ***Time out (s)*** | ***Latent hold (s)*** | ***ITI (s)*** | ***Criteria*** |
| --- | --- | --- | --- | --- | --- | --- | --- | --- |
| *1* | Collect pellets | 100 | 20 | - | - | - | - | ≥80 trials, 1 day |
| *2* | Nose poke training | 100 | 30 | - | - | - | 0, 2, 4, 6, 8, 10 | ≥80 trials, 2 days |
| *3* | Signal detection training | 120 | 30 | Unlimited | - | - | 1, 2, 3 | ≥80% Correct, 2 days |
| *4* | Signal detection | 120 | 30 | 1 | 5 | 4 | 1, 2, 3 | ≥80% Correct, 2 days |
| *Final SDT* | SDT with vSD | 120 | 30 | 0.03; 0.06; 0.12; 0.25; 1* | 5 | 4 | 5 | Stable baseline (>15 sessions) |

**Table S2**: Estimated marginal means (emmeans) and standard error of the difference (SED) for each AMPH dose for the whole cohort and attentional subcategories. Bold numbers indicate a statistically significant difference, whereas underlined numbers indicate a trend towards significance (p<0.11) between vehicle and the respective dose for that category in the *post-hoc* test. There was no significant dose x performance group x SD interaction; F_(30, 529)_ = 0.76, p = n.s.

|  | Amphetamine | | | | | | | | | | | | | | | |
| --- | --- | --- | --- | --- | --- | --- | --- | --- | --- | --- | --- | --- | --- | --- | --- | --- |
|  | Veh | | | | 0.1 mg/kg | | | | 0.2 mg/kg | | | | 0.4 mg/kg | | | |
|  | All | LA | MA | HA | All | LA | MA | HA | All | LA | MA | HA | All | LA | MA | HA |
| **Accuracy** | 78.3 ±2.2 | 73.9  ±2.2 | 77.2  ±2.1 | 83.4  ±2.1 | 78.6 ±2.2 | 75.2  ±2.2 | 80.8  ±2.1 | 79.7  ±2.1 | 77.9 ±2.2 | 76.4  ±2.2 | 75.2  ±2.2 | 82.3  ±2.1 | 78.4 ±2.2 | 75.2  ±2.2 | 76.4  ±2.2 | 83.6  ±2.1 |
| **d’** | 1.40 ±0.2 | 0.96 ±0.2 | 1.32  ±0.2 | 1.91 ±0.2 | 1.44 ±0.2 | 1.14 ±0.2 | 1.71  ±0.2 | 1.46 ±0.2 | 1.37 ±0.2 | 1.15 ±0.2 | 1.19  ±0.2 | 1.77 ±0.2 | 1.50 ±0.2 | 1.06 ±0.2 | 1.47  ±0.2 | 1.96 ±0.2 |
| **β** | 0.80 ±0.1 | 0.67 ±0.1 | 0.82  ±0.1 | 0.89 ±0.1 | 0.73 ±0.1 | 0.70 ±0.1 | 0.81  ±0.1 | 0.67 ±0.1 | 0.77 ±0.1 | 0.63 ±0.1 | 0.84  ±0.1 | 0.82 ±0.1 | 0.92 ±0.1 | 0.69 ±0.1 | **1.14**  **±0.1** | 0.91 ±0.1 |
| ***v*** | 17.8 ±2.4 | 16.4 ±3.8 | 15.3  ±3.5 | 21.6 ±3.5 | 20.5 ±2.4 | 16.5 ±3.8 | 24.7  ±3.5 | 20.0 ±3.5 | 18.3 ±2.4 | 17.8 ±3.8 | 14.3  ±3.5 | 22.7 ±3.5 | 18.3 ±2.4 | 14.4 ±3.8 | 15.5  ±3.5 | 24.6 ±3.5 |
| ***p*_s_** | 0.36 ±0.06 | 0.48 ±0.05 | 0.35  ±0.05 | 0.24 ±0.05 | 0.35 ±0.06 | 0.42 ±0.05 | 0.27  ±0.05 | 0.36 ±0.05 | 0.37 ±0.06 | 0.49 ±0.05 | 0.37  ±0.05 | 0.26 ±0.05 | 0.33 ±0.06 | 0.48 ±0.05 | 0.30  ±0.05 | 0.20 ±0.05 |
| **No signal** | 68.4 ±6.0 | 55.4 ±4.3 | 70.9  ±4.1 | 78.4  ±4.1 | 68.6 ±6.0 | 61.4 ±4.3 | 72.8  ±4.1 | 71.9  ±4.1 | 69.8 ±6.0 | 57.9 ±4.3 | 70.6  ±4.1 | 80.6  ±4.1 | 72.4 ±6.0 | 58.9 ±4.3 | 77.8  ±4.1 | 80.0  ±4.1 |
| **30 ms** | 61.7 ±3.5 | 60.9 ±6.3 | 58.4  ±5.9 | 65.8 ±5.9 | 60.3 ±3.5 | 56.2 ±6.3 | 60.0  ±5.9 | 64.2 ±5.9 | 59.1 ±3.5 | 65.7 ±6.3 | 54.1  ±5.9 | 58.3 ±5.9 | 55.1 ±3.5 | 60.9 ±6.3 | 47.5  ±5.9 | 57.5 ±5.9 |
| **60 ms** | 81.2 ±2.5 | 80.0 ±4.0 | 78.3  ±3.7 | 85.0 ±3.7 | 84.0 ±2.5 | 82.9 ±4.0 | 85.0  ±3.7 | 84.2 ±3.7 | 85.2 ±2.5 | 86.7 ±4.0 | 79.2  ±3.7 | 90.0 ±3.7 | 82.3 ±2.5 | 81.0 ±4.0 | 78.3  ±3.7 | 87.5 ±3.7 |
| **120 ms** | 92.2 ±2.2 | 91.4 ±3.4 | 91.7  ±3.2 | 93.3 ±2.2 | 91.0 ±2.2 | 91.4 ±3.4 | 89.2  ±3.2 | 92.5 ±3.2 | 90.7 ±2.2 | 89.5 ±3.4 | 86.7  ±3.2 | 95.8 ±3.2 | 92.4 ±2.2 | 92.4 ±3.4 | 88.3  ±3.2 | 96.7 ±3.2 |
| **250 ms** | 95.1 ±1.3 | 95.2 ±2.3 | 95.0  ±2.2 | 95.0 ±2.2 | 96.8 ±1.3 | 92.4 ±2.3 | 99.2  ±2.2 | 98.3 ±2.2 | 95.6 ±1.3 | 96.2 ±2.3 | 96.7  ±2.2 | 94.2 ±2.2 | 95.6 ±1.3 | 93.3 ±2.3 | 95.8  ±2.2 | 97.5 ±2.2 |
| **1 s** | 96.1 ±1.3 | 92.1 ±2.0 | 97.5  ±1.8 | 98.1 ±1.8 | 96.7 ±1.3 | 95.7 ±2.0 | 96.2  ±1.8 | 98.1 ±1.8 | 95.4 ±1.3 | 91.4 ±2.0 | 95.6  ±1.8 | 98.8 ±1.8 | 97.6 ±1.3 | **98.6 ±2.0** | 97.5  ±1.8 | 96.9 ±1.8 |
| **Trial initiation latency** | 0.91 ±0.1 | 1.06 ±0.1 | 0.86  ±0.1 | 0.82 ±0.1 | 0.86 ±0.1 | 1.03 ±0.1 | 0.84  ±0.1 | 0.73 ±0.1 | **0.82 ±0.1** | 0.99 ±0.1 | 0.79  ±0.1 | 0.68 ±0.1 | **0.73 ±0.1** | **0.84 ±0.1** | 0.72  ±0.1 | 0.64 ±0.1 |
| **Response latency** | 1.21 ±0.01 | 1.19 ±0.01 | 1.22  ±0.01 | 1.21 ±0.01 | 1.21 ±0.01 | 1.21 ±0.01 | 1.21  ±0.01 | 1.22 ±0.01 | 1.20 ±0.01 | 1.20 ±0.01 | 1.20  ±0.01 | 1.21 ±0.01 | 1.21 ±0.01 | 1.21 ±0.01 | 1.21  ±0.01 | 1.20 ±0.01 |
| **Anticipatory responses** | 170 ±9.6 | 176 ±13.1 | 148  ±12.2 | 185 ±12.2 | 179 ±9.6 | 183 ±13.1 | 162  ±12.2 | 192 ±12.2 | **186 ±9.6** | 199 ±13.1 | **168**  **±12.2** | 193 ±12.2 | **183 ±9.6** | 189 ±13.1 | **168**  **±12.2** | 192 ±12.2 |

**Table S3**: Estimated marginal means (emmeans) and standard error of the difference (SED) for each MPH dose for the whole cohort as well as the low and high attentive subcategories. Bold numbers indicate a statistically significant difference, whereas underlined numbers indicate a trend towards significance (p<0.11) between vehicle and the respective dose for that category in the *post-hoc* test. There was no significant dose x performance group x SD interaction; F_(30, 529)_ = 0.51, p = n.s.

|  | Methylphenidate | | | | | | | | | | | | | | | |
| --- | --- | --- | --- | --- | --- | --- | --- | --- | --- | --- | --- | --- | --- | --- | --- | --- |
|  | Veh | | | | 0.3 mg/kg | | | | 1 mg/kg | | | | 3 mg/kg | | | |
|  | All | LA | MA | HA | All | LA | MA | HA | All | LA | MA | HA | All | LA | MA | HA |
| **Accuracy** | 79.2 ±3.1 | 74.8  ±1.9 | 77.0  ±1.8 | 85.6  ±1.8 | 79.8 ±3.1 | 75.9  ±1.9 | 77.7  ±1.8 | 85.9  ±1.8 | 79.7 ±3.1 | 74.6  ±1.9 | 78.8  ±1.8 | 85.6  ±1.8 | 77.9 ±3.1 | 75.4  ±1.9 | 75.3  ±1.8 | 83.1  ±1.8 |
| **d’** | 1.52 ±0.3 | 1.14 ±0.2 | 1.39  ±0.2 | 2.03 ±0.2 | 1.65 ±0.3 | 1.28 ±0.2 | 1.48  ±0.2 | 2.18 ±0.2 | 1.55±0.3 | 1.07 ±0.2 | 1.57  ±0.2 | 2.00 ±0.2 | 1.39 ±0.3 | 1.13 ±0.2 | 1.25  ±0.2 | 1.80 ±0.2 |
| **β** | 0.88 ±0.1 | 0.75 ±0.2 | 0.96  ±0.2 | 0.93 ±0.2 | 1.01 ±0.1 | 0.82 ±0.2 | 1.15  ±0.2 | 1.05 ±0.2 | 0.73 ±0.1 | 0.76 ±0.2 | 0.89  ±0.2 | 0.54 ±0.2 | 0.84 ±0.1 | 0.77 ±0.2 | 0.98  ±0.2 | 0.75 ±0.2 |
| ***v*** | 20.9 ±5.2 | 13.6 ±3.9 | 17.6  ±3.7 | 31.5 ±3.5 | 21.5 ±5.2 | 17.5 ±3.9 | 18.0  ±3.7 | 29.4 ±3.7 | 24.1 ±5.2 | 14.9 ±3.9 | 21.3  ±3.7 | 35.7 ±3.7 | 18.3 ±5.2 | 13.0 ±3.9 | 14.9  ±3.7 | 27.0 ±3.7 |
| ***p*_s_** | 0.29 ±0.05 | 0.40 ±0.06 | 0.29  ±0.05 | 0.18 ±0.05 | 0.27 ±0.05 | 0.32 ±0.06 | 0.31  ±.5 | 0.17 ±0.05 | 0.32 ±0.05 | 0.42 ±0.06 | 0.25  ±0.05 | 0.29 ±0.05 | 0.36 ±0.05 | 0.43 ±0.06 | 0.37  ±0.05 | 0.28 ±0.05 |
| **No signal** | 73.4 ±4.4 | 67.5  ±4.2 | 73.4  ±4.0 | 79.4  ±4.0 | 78.0 ±4.4 | 69.6  ±4.2 | 79.4  ±4.0 | 84.7  ±4.0 | 73.9 ±4.4 | 63.9  ±4.2 | 75.9  ±4.0 | 81.2  ±4.0 | 70.6 ±4.4 | 63.9  ±4.2 | 70.3  ±4.0 | 77.5  ±4.0 |
| **30 ms** | 55.0 ±5.0 | 57.1 ±6.2 | 45.0  ±5.8 | 63.3 ±5.8 | 57.9 ±5.0 | 54.3 ±6.2 | 52.5  ±5.8 | 66.7 ±5.8 | 60.5 ±5.0 | 57.2 ±6.2 | 50.8  ±5.8 | 73.3 ±5.8 | **63.2 ±5.0** | 62.9 ±6.2 | **60.0**  **±5.8** | 66.7 ±5.8 |
| **60 ms** | 83.7 ±3.7 | 79.0 ±4.5 | 81.7  ±4.3 | 90.0 ±4.3 | 79.6 ±3.7 | 76.2 ±4.5 | 79.2  ±4.3 | 83.3 ±4.3 | 82.8 ±3.7 | 79.0 ±4.5 | 78.3  ±4.3 | 90.8 ±4.3 | 78.4 ±3.7 | 76.2 ±4.5 | 73.3  ±4.3 | 85.8 ±4.3 |
| **120 ms** | 92.3 ±2.7 | 86.7 ±2.7 | 94.2  ±2.5 | 95.8 ±2.5 | 92.9 ±2.7 | 87.6 ±2.7 | 93.3  ±2.5 | 97.5 ±2.5 | 93.5 ±2.7 | 89.5 ±2.7 | 93.3  ±2.5 | 97.5 ±2.5 | 91.1 ±2.7 | 88.6 ±2.7 | 89.2  ±2.5 | 95.8 ±2.5 |
| **250 ms** | 97.1 ±0.9 | 94.3 ±1.6 | 98.3  ±1.5 | 98.3 ±1.5 | 98.5 ±0.9 | **99.0 ±1.6** | 97.5  ±1.5 | 99.2 ±1.5 | 98.0 ±0.9 | 98.1 ±1.6 | 99.2  ±1.5 | 96.7 ±1.5 | 96.5 ±0.9 | 96.2 ±1.6 | 95.0  ±1.5 | 98.3 ±1.5 |
| **1 s** | 97.1 ±1.2 | 97.1 ±1.8 | 97.5  ±1.7 | 96.9 ±1.7 | 97.1 ±1.2 | 96.4 ±1.8 | 96.9  ±1.7 | 98.1 ±1.7 | 96.5 ±1.2 | 94.3 ±1.8 | 96.9  ±1.7 | 98.1 ±1.7 | 65.6 ±1.2 | 92.1 ±1.8 | 95.6  ±1.7 | 98.8 ±1.7 |
| **Trial initiation latency** | 0.80 ±0.1 | 1.01 ±0.1 | 0.73  ±0.1 | 0.67 ±0.1 | 0.76 ±0.1 | 0.89 ±0.1 | 0.75  ±0.1 | 0.65 ±0.1 | 0.77 ±0.1 | 1.06 ±0.1 | 0.65  ±0.1 | 0.62 ±0.1 | 0.71 ±0.1 | 0.81 ±0.1 | 0.66  ±0.1 | 0.66 ±0.1 |
| **Response latency** | 1.21 ±0.01 | 1.19 ±0.01 | 1.22  ±0.01 | 1.21 ±0.01 | 1.21 ±0.01 | 1.20 ±0.01 | 1.22  ±0.01 | 1.21 ±0.01 | 1.20 ±0.01 | 1.20 ±0.01 | 1.20  ±0.01 | 1.20 ±0.01 | 1.21 ±0.01 | 1.21 ±0.01 | 1.22  ±0.01 | 1.19 ±0.01 |
| **Anticipatory responses** | 179 ±9.6 | 190 ±13.3 | 159  ±12.5 | 190 ±12.2 | 173 ±9.6 | 184 ±13.3 | 158  ±12.5 | 179 ±12.2 | 184 ±9.6 | 190 ±13.3 | 166  ±12.5 | 195 ±12.2 | 192 ±9.6 | 191 ±13.3 | 178  ±12.5 | 206 ±12.2 |

**Table S4**: Estimated marginal means (emmeans) and standard error of the difference (SED) for each ATO dose for the whole cohort as well as the low and high attentive subcategories. Bold numbers indicate a statistically significant difference, whereas underlined numbers indicate a trend towards significance (p<0.11) between vehicle and the respective dose for that category in the *post-hoc* test. There was no significant dose x performance group x SD interaction; F_(30, 552)_ = 0.70, p = n.s.

|  | Atomoxetine | | | | | | | | | | | | | | | |
| --- | --- | --- | --- | --- | --- | --- | --- | --- | --- | --- | --- | --- | --- | --- | --- | --- |
|  | Veh | | | | 0.1 mg/kg | | | | 0.3 mg/kg | | | | 1 mg/kg | | | |
|  | All | LA | MA | HA | All | LA | MA | HA | All | LA | MA | HA | All | LA | MA | HA |
| **Accuracy** | 82.9 ±3.2 | 80.2  ±1.9 | 82.3  ±1.9 | 86.2  ±1.9 | **79.1 ±3.2** | **73.1**  **±1.9** | 78.4  ±1.9 | 85.8  ±1.9 | **79.4 ±3.2** | **70.9**  **±1.9** | 80.9  ±1.9 | 86.2  ±1.9 | **78.6 ±3.2** | **74.7**  **±1.9** | 77.3  ±1.9 | 83.8  ±1.9 |
| **d’** | 1.85 ±0.3 | 1.60 ±0.2 | 1.80  ±0.2 | 2.15 ±0.2 | 1.62 ±0.3 | **1.13 ±0.2** | 1.57  ±0.2 | 2.18 ±0.2 | 1.62 ±0.3 | **0.88 ±0.2** | 1.76  ±0.2 | 2.21 ±0.2 | **1.53 ±0.3** | **1.18 ±0.2** | 1.50  ±0.2 | 1.92 ±0.2 |
| **β** | 0.81 ±0.1 | 0.83 ±0.2 | 0.76  ±0.2 | 0.84 ±0.2 | 1.02 ±0.1 | 0.91 ±0.2 | 1.00  ±0.2 | 1.15 ±0.2 | 1.00 ±0.1 | 0.95 ±0.2 | 1.13  ±0.2 | 0.93 ±0.2 | 1.02 ±0.1 | 1.02 ±0.2 | 1.34  ±0.2 | 0.90 ±0.2 |
| ***v*** | 25.6 ±3.9 | 22.1 ±3.4 | 25.8  ±3.4 | 29.0 ±3.4 | **19.7 ±3.9** | **13.6 ±3.4** | **17.7**  **±3.4** | 27.8 ±3.4 | 21.9 ±3.9 | **10.7 ±3.4** | 24.6  ±3.4 | 30.4 ±3.4 | **19.4 ±3.9** | 15.6 ±3.8 | **16.7**  **±3.4** | 25.8 ±3.5 |
| ***p*_s_** | 0.26 ±0.05 | 0.30 ±0.05 | 0.29  ±0.05 | 0.20 ±0.05 | 0.27 ±0.05 | 0.32 ±0.05 | 0.27  ±0.05 | 0.20 ±0.05 | 0.29 ±0.05 | 0.43 ±0.05 | 0.24  ±0.05 | 0.20 ±0.05 | 0.30 ±0.05 | 0.37 ±0.05 | 0.29  ±0.05 | 0.24 ±0.05 |
| **No signal** | 76.7 ±4.3 | 74.4  ±3.7 | 74.1  ±3.7 | 81.6  ±3.7 | 75.5 ±4.3 | 67.8  ±3.7 | 74.7  ±3.7 | 84.1  ±3.7 | 75.9 ±4.3 | 67.2  ±3.7 | 73.8  ±3.7 | 86.9  ±3.7 | 76.0 ±4.3 | 69.1  ±3.7 | 77.5  ±3.7 | 81.6  ±3.7 |
| **30 ms** | 64.4 ±4.8 | 58.3 ±6.9 | 65.8  ±6.9 | 69.2 ±6.9 | **53.1 ±4.8** | 42.5 ±6.9 | 51.7  ±6.9 | 65.0 ±6.9 | 58.6 ±4.8 | 55.0 ±6.9 | 52.5  ±6.9 | 68.3 ±6.9 | **55.3 ±4.8** | 54.2 ±6.9 | 50.8  ±6.9 | 60.8 ±6.9 |
| **60 ms** | 86.9 ±4.0 | 83.3 ±4.3 | 88.3  ±4.3 | 89.2 ±4.3 | 82.8 ±4.0 | 75.8 ±4.3 | 83.3  ±4.3 | 89.1 ±4.3 | 80.8 ±4.0 | **67.5 ±4.3** | 87.5  ±4.3 | 87.5 ±4.3 | 82.2 ±4.0 | 79.2 ±4.3 | 79.2  ±4.3 | 88.3 ±4.3 |
| **120 ms** | 93.9 ±2.2 | 95.0 ±2.3 | 91.7  ±2.3 | 95.0 ±2.3 | 91.1 ±2.2 | **88.3 ±2.3** | 88.3  ±2.3 | 96.7 ±2.3 | 93.3 ±2.2 | 90.0 ±2.3 | 93.3  ±2.3 | 96.7 ±2.3 | 91.9 ±2.2 | **86.7 ±2.3** | 91.7  ±2.3 | 97.5 ±2.3 |
| **250 ms** | 96.9 ±1.0 | 95.8 ±1.8 | 97.5  ±1.8 | 97.5 ±1.8 | 96.4 ±1.0 | 95.8 ±1.8 | 95.8  ±1.8 | 97.5 ±1.8 | 96.7 ±1.0 | 93.3 ±1.8 | 99.2  ±1.8 | 97.5 ±1.8 | 96.7 ±1.0 | 97.5 ±1.8 | 95.8  ±1.8 | 96.7 ±1.8 |
| **1 s** | 97.3 ±1.2 | 96.9 ±1.9 | 96.2  ±1.9 | 98.8 ±1.9 | 96.5 ±1.2 | 93.8 ±1.9 | 96.9  ±1.9 | 98.8 ±1.9 | 96.2 ±1.2 | 96.2 ±1.9 | 94.4  ±1.9 | 98.1 ±1.9 | 96.9 ±1.2 | 96.9 ±1.9 | 95.6  ±1.9 | 98.1 ±1.9 |
| **Trial initiation latency** | 0.74 ±0.1 | 0.74 ±0.1 | 0.80  ±0.1 | 0.66 ±0.1 | 0.70 ±0.1 | 0.71 ±0.1 | 0.80  ±0.1 | 0.59 ±0.1 | 0.74 ±0.1 | 0.80 ±0.1 | 0.76  ±0.1 | 0.66 ±0.1 | 0.77 ±0.1 | 0.81 ±0.1 | 0.83  ±0.1 | 0.68 ±0.1 |
| **Response latency** | 1.19 ±0.01 | 1.17 ±0.02 | 1.19  ±0.02 | 1.21 ±0.02 | 1.21 ±0.01 | 1.20 ±0.02 | 1.21  ±0.02 | 1.21 ±0.02 | 1.20 ±0.01 | 1.19 ±0.02 | 1.20  ±0.02 | 1.21 ±0.02 | 1.20 ±0.01 | 1.19 ±0.02 | 1.21  ±0.02 | 1.20 ±0.02 |
| **Anticipatory responses** | 170 ±9.0 | 158 ±15.8 | 174  ±15.8 | 179 ±15.8 | 169 ±9.0 | 153 ±15.8 | 180  ±15.8 | 175 ±15.8 | 169 ±9.0 | 154 ±15.8 | 176  ±15.8 | 177 ±15.8 | **150 ±9.0** | **132 ±15.8** | 164  ±15.8 | **153 ±15.8** |
